# Pasture age impacts soil fungal composition while bacteria respond to soil chemistry

**DOI:** 10.1101/2021.09.13.460094

**Authors:** Fiona M. Seaton, Robert I. Griffiths, Tim Goodall, Inma Lebron, Lisa R. Norton

## Abstract

Pasture is a globally important managed habitat providing both food and income. The way in which it is managed leads to a wide range of impacts on soil microbial communities and associated soil health. While there have been several studies comparing pasture farming to other forms of land use, we still have limited understanding of how the soil microbial communities vary between pasture farms and according to management practices. Here we present the results of a field survey across 56 UK livestock farms that are managed by members of the Pasture fed Livestock Association, using amplicon sequencing of the 16S and ITS regions to characterise the soil bacterial and fungal community within fields that have been under pasture for differing durations. We show that grazing management intensity has only limited effects upon microbial community structure, while the duration of pasture since ploughing (ranging from 1 year to over 100 years) impacted the fungal community structure. The impact of management duration was conditional upon soil physicochemical properties, particularly pH. Plant community effects on upon soil bacterial and fungal composition appear to also interact with the soil chemistry, highlighting the importance of plant-soil interactions in determining microbial community structure. Analyses of microbial indicators revealed proportionally more fungal taxa that responded to multiple ecosystem health associated properties than bacterial taxa. We also identified several fungal taxa that both acted as indicators of soil health related properties within our dataset and showed differentiation between grassland types in a national survey, indicating the generality of some fungal indicators to the national level. Members of the Agaricomycetes were associated with multiple indicators of soil health. Our results show the importance of maintaining grassland for the development of plant-soil interactions and microbial community structure with concomitant effects on soil and general ecosystem health.

**Highlights:**

- Soil bacteria and fungi from >50 pastures were analysed using DNA metabarcoding
- Duration of pasture since ploughing impacted fungal community composition
- Bacteria were more associated with soil physicochemical properties than fungi
- Plant effects on soil microbes were through interactions with soil chemistry
- Microbial indicators of soil health were disproportionately fungal taxa

## Introduction

How we manage our soil health is of increasing importance, as we balance increasing food demand with the need to maintain soil lifespans and simultaneously provide valuable ecosystem services such as carbon storage (Bossio et al., 2020; Evans et al., 2020). Livestock farming takes up large areas of land and contributes approximately 15% of anthropogenic greenhouse gas emissions, however it is globally important for both nutrition and income security (Gerber et al., 2013; Smith et al., 2013). Balancing the demands of livestock farming while maintaining and promoting ecosystem health to ensure production of both food and ecosystem services from these systems for future generations requires innovative farming practices coupled with whole-system evaluation of the effects of these different farm managements upon ecosystem and soil health (Kibblewhite et al., 2008). Soil biodiversity is a key component of soil health, as it underpins a variety of ecosystem functions (Bardgett & van der Putten, 2014; Delgado-Baquerizo et al., 2020). Land use intensity is known to influence soil ecological communities, with more intense land use types leading to reductions in diversity and biomass and changes in microbially-mediated soil functions (Neal et al., 2021; Tsiafouli et al., 2015). Soil microbial communities also respond to land use, however unlike soil animals they can show less consistent changes in diversity and composition between differing types of grassland and arable systems (George et al., 2019).

Pasture systems have been found to have greater microbial biomass, enzyme activities and differing microbial compositions to crop systems (Acosta-Martínez et al., 2010; Chen et al., 2018; Walkup et al., 2020). In comparison to forests, however, pasture systems can show lower biomass and functional diversity, which has been related to both differences in vegetation diversity and plant litter inputs (Cardozo Junior et al., 2018; Chen et al., 2018; Silveira Sartori Silva et al., 2019). Many of these differences in microbial communities’ responses to land use may be related to changes in soil physicochemical properties, for example soil pH is known to be a strong driver of microbial community diversity, composition and activities (George et al., 2019; Griffiths et al., 2011; Hendershot et al., 2017). Lauber et al. (2008) found that changes in soil pH and C:N ratios explained the variation in bacterial and fungal communities respectively across pasture, crop and forest land uses. Soil structural properties are also known to both influence and be influenced by microbial activities, e.g. the microbial contribution to aggregate formation, stabilisation and eventually degradation has been extensively reviewed (Lynch & Bragg, 1985; Oades, 1993; Totsche et al., 2010). However, we are still starting to unfold the mechanisms underlying the interacting microbial, chemical and structural responses to land use practices within differing environmental contexts, and how much of this information is transferable to inferring responses within land use types to management changes.

Certain management decisions made within pasture farming have been found to have an impact upon soil microbial communities. In particular, fertilisation of grasslands can alter soil microbial composition and biomass (Leff et al., 2015; Walkup et al., 2020). An observational study of pasture types across New Zealand also found that dairy pasture shows distinct bacterial communities compared to other types of pasture use (Dignam et al., 2018). The composition of the plant community in pasture systems has also been found to influence soil fungal community composition (Zheng et al., 2016), although plant community composition was found to have limited impact upon microbial biomass and activities across a variety of Brazilian pasture systems (Cardozo Junior et al., 2018).

The Pasture fed Livestock Association (PFLA) was set up in 2009 by a group of British farmers who wished to promote the human health benefits of purely grass-fed cattle products, and who created a set of certification standards for pasture-fed ruminant livestock (Pasture for Life Association, 2020; Vetter, 2020). While the initial motivations of the Association were related to the potential health benefits of grass-fed beef and dairy products (Daley et al., 2010; Haskins et al., 2019), the PFLA standards also include provisions for the protection of wildlife, reduction of inputs and maintenance of general ecosystem health. These considerations are particularly important due to the higher land use and thus potentially higher greenhouse gas emissions associated with grass-fed cattle. However, this may be counteracted by the tendency for grass-fed cattle to be reared on land unsuitable for crops and potential gains in biodiversity and other ecosystem-related functions, compared to more intensive production systems including grain-fed cattle or arable systems (Clark & Tilman, 2017). It will also depend on levels of inputs, such as mineral fertilisers and pesticides which are very low on PFLA farms. Within this work we examine the impact of pasture management intensity and duration upon soil microbial communities through analysis of 56 PFLA farms across Great Britain. We combine bacterial and fungal community composition measurements from DNA sequencing with detailed farm management data from farmer surveys, soil physicochemical measurements and plant surveys. We compare our microbial data to microbial data taken from a variety of grassland types across the British landscape within the Countryside Survey of Great Britain to evaluate if there are common indicators of grassland type relating to management (Carey et al., 2008). Our hypotheses were:

1. Soil bacterial and fungal communities under PFLA management relate to both current management and the duration of these management practices
2. Plant composition impacts on soil microbial composition through changes in key plant species cover (e.g. *Lolium perenne*)
3. Taxa that responded to management practices and soil health indicators on the PFLA farms would also show differentiation between grassland types in the broader dataset

## Methods

### Field survey

In total 56 PFLA member farms were surveyed across Great Britain, in the summer of 2018 (May to early September). Soil and vegetation were sampled using protocols from the Countryside Survey (CS), see Emmett et al. (2008); Maskell et al. (2008); and Wood et al. (2017) to enable comparison with data collected as part of the survey in 2007. Sampling took place within a single field with the exception of one farm on which data was collected from two fields of differing land use history, hence n=57. The sampling location was determined pre-survey and validated with the farmer on site (to check that an atypical field had not been selected), and included a 200m^2^ plot surveyed for vegetation composition and a soil core taken from a central point within the square. Farmers were interviewed to collect detailed management data about the current and historical uses of the field, and to ascertain other relevant variables including age of pasture since plough, type of grazing, fertilisation regime and organic status. Plots were allocated to grassland Broad Habitat types, hereafter referred to as ‘grassland types’ (Improved, Neutral, Acid and Calcareous) according to the CS field habitat key by surveyors in the field (Jackson, 2000; Maskell et al., 2008). Plant Ellenberg scores were calculated for each plot based on averages of the different Ellenberg scores as identified for each plant species present within PLANTATT (Hill et al., 2004).

### Soil physicochemical properties

Soil physicochemical properties were measured following the methods of Emmett et al. (2008). In brief, a single soil core (15cm depth and 7cm diameter) was taken from for each plot and tested for soil organic carbon, total carbon, total nitrogen, pH, Olsen P, total P, electrical conductivity, bulk density, aggregate stability, clay, silt and sand content. Soil pH and electrical conductivity were measured in a 1:2.5 weight suspension in deionised water with a pH meter (Corning 220) and a conductivity meter (Jenway 4510) respectively and pH was also measured in a 0.01 M CaCl_2_ solution. Soil organic carbon was measured by loss on ignition, while total soil carbon and soil nitrogen were measured by an Elementar Vario-EL elemental analyser (Elementaranalysensysteme GmbH, Hanau, Germany). Soil phosphorus is measured both as total P and using the Olsen-P method. Soil texture was measured by laser granulometry with the Beckman Coulter LS 13 320 as described in Seaton et al. (2020). Aggregate stability was measured using wet sieving with a Eijkelkamp apparatus, where 2g of aggregates were immersed in deionised water within a sieve of 250μm aperture and agitated for half an hour then dried and weighed (weight A). The remaining aggregates are then immersed in NaOH and agitated for another half hour, then dried and weighed again (weight B). Aggregate stability is calculated as (weight A – weight B)/initial weight.

### Soil microbial community characterisation

DNA was extracted from 0.2 g of soil using the PowerSoil-htp 96 Well DNA Isolation kit (Qiagen, Hiden, Germany) according to manufacturer’s protocols. For bacterial 16S rRNA amplicons the dual indexing protocol of Kozich et al. (2013) was used for MiSeq sequencing (Illumina, San Diego, US) with each primer consisting of the appropriate Illumina adapter, 8-nt index sequence, a 10-nt pad sequence, a 2-nt linker and the amplicon specific primer. The V3–V4 hypervariable regions of the bacterial 16S rRNA gene were amplified using primers 341F (Muyzer et al., 1993) and 806R (Yu et al., 2005), CCTACGGGAGGCAGCAG and GCTATTGGAGCTGGAATTAC respectively. Amplicons were generated using a high-fidelity DNA polymerase Q5 Taq (New England Biolabs, Ipswich, US). After an initial denaturation at 95 °C for 2 minutes PCR conditions were: denaturation at 95 °C for 15 seconds; annealing at 55 °C for 30 seconds with extension at 72 °C for 30 seconds, repeated for 30 cycles; final extension of 10 minutes at 72 °C was included.

Fungal internal transcribed spacer (ITS) amplicon sequences were generated using a 2-step amplification approach. Primers GTGARTCATCGAATCTTTG and TCCTCCGCTTATTGATATGC (Ihrmark et al., 2012) were each modified at 5’ end with the addition of Illumina pre-adapter and Nextera sequencing primer sequences. After an initial denaturation at 95°C for 2 minutes, PCR conditions were: denaturation at 95°C for 15 seconds; annealing at 52°C for 30 seconds with extension at 72°C for 30 seconds; repeated for 25 cycles. A final extension of 10 minutes at 72°C was included.

PCR products were cleaned using a ZR-96 DNA Clean-up Kit (Zymo Research Inc., Irvine, US) following manufacturer’s instructions. MiSeq adapters and 8nt dual-indexing barcode sequences were added during a second step of PCR amplification. After an initial denaturation 95°C for 2 minutes, PCR conditions were: denaturation at 95°C for 15 seconds; annealing at 55°C for 30 seconds with extension at 72°C for 30 seconds; repeated for 8 cycles with a final extension of 10 minutes at 72°C.

Amplicon concentrations were normalized using SequalPrep Normalization Plate Kit (Thermo Fisher Scientific, Waltham, US) and amplicon sizes determined using an 2200 TapeStation (Agilent, Santa Clara, US) prior to sequencing each amplicon library separately using MiSeq (Illumina, San Diego, US) with V3 600 cycle reagents at concentrations of 6 and 12 pM (16S and ITS respectively) with a 5% PhiX control library.

Illumina demultiplexed sequences for 16S and ITS are available at the European Nucleotide Archive under PRJEB46195 primary accession code, sample accession codes ERS7103229 to ERS7103287. Previously generated amplicon data (generated utilising the same amplicon primers and DNA extraction methodology as this study) from the GB Countryside Survey of 2007 (data will be made available on the ENA by acceptance), was bioinformatically processed in parallel, allowing comparison of this sample set to a national survey. Each amplicon and sequencing run were processed separately in R using DADA2 (Callahan et al., 2016), with a Cutadapt (Martin, 2011) step added for ITS sequences. Forward 16S amplicon reads were truncated to 250-nt. Sequences with Ns and error greater than maxEE=1 were removed. Sequences were dereplicated and the DADA2 core sequence variant inference algorithm applied and actual sequence variants (ASVs) generated. Chimeric sequences were removed using removeBimeraDenovo at default settings. ASVs were subject to taxonomic assignment using assignTaxonomy at default settings with GreenGenes v13.8 (DeSantis et al., 2006; McDonald et al., 2012) training database.

ITS amplicon reads we pre-processed using Cutadapt to remove primer sequences and negate read-through risk. Using DADA2 reads were then truncated to 205-nt and 160-nt (forward and reverse, respectively). Sequences with Ns and error greater than maxEE=5 were removed. Using default settings sequences were dereplicated into ASVs, denoised, merged, chimera checked and assigned taxonomies using Unite v7.2 (Kõljalg et al., 2005) training database.

Fungal sequences were matched against funguild to get information on trophic modes (Nguyen et al., 2016). ASVs that occurred in negative controls were removed from the results and the sequence tables were limited to ASVs identified as bacteria, minus chloroplast and mitochondrial ASVs, and fungi respectively.

### Statistical analysis

All samples were rarefied to 22000 reads for bacteria and 5000 reads for fungi, resulting in keeping all 57 samples. Rarefaction was repeated 10 times and the average result taken for richness, Shannon diversity, Simpson diversity and the average occurrence matrix. The impact of pasture age (an ordered factor) upon microbial richness, LOI, Olsen P and *L*. *perenne* cover was tested by fitting Bayesian ordinal regression models in the brms package (Bürkner, 2017; Bürkner & Vuorre, 2019).

The impact of field management and plant and soil properties upon the microbial composition was explored using Bray-Curtis distance to summarise the rarefied communities (McKnight et al., 2019). This included NMDS with the associated envfit function for numerical variables, multivariate homogeneity of group dispersions using the betadisper function and distance-based redundancy analysis (dbRDA) using the capscale function with 99999 permutations all in the vegan package version 2.5-7 (Oksanen et al., 2020).

Variation partitioning was performed on the NMDS scores when done for 4 dimensions, with the soil structural metrics, soil chemical data, plant community composition, and the farm management data as predictors. Soil structural data included aggregate stability, bulk density, sand, silt and clay. Sand, silt and clay were all converted using a centred log ratio transformation and then the first 4 scores from principal component analysis (PCA) on the centred, scaled data used in varpart. Soil chemical data included pH in water, conductivity, total carbon, total nitrogen, Olsen P and was also represented using the first 4 axis scores of PCA. The plants identified at each plot and their total cover was used to construct Bray-Curtis distances for a principal coordinates analysis (PCoA) which was run using the ape package (Paradis & Schliep, 2019). Select categorical farm management data properties were included in the variation partitioning from the axis scores from a 4-dimensional multiple correspondence analysis (MCA) built using the MASS package (Venables & Ripley, 2002).The farm management data included in the MCA were: permanent or temporary pasture; age of pasture since plough (<20, 20-50 and >50 years); type of grazing (mob, set stocking, or paddock/strip); organic status; winter grazing (yes/no/sometimes); fertiliser use (mineral with liming, mineral only or manure); sheep presence.

The DeSeq2 package (Love et al., 2014) was used on the unrarefied data to identify indicator taxa of field age (using cutoffs of 20 year or 50 years), organic carbon (LOI), Olsen-P and Lolium perenne cover. Each of these variables were fit to the data in a model including pH and then only the taxa that responded to the variable of interest at p < 0.05 extracted. All analysis was done in R version 4.0.2 (R Core Team, 2020).

## Results

In total, 31340 bacterial ASVs were recovered across all samples, 9 archaeal ASVs and 6756 fungal ASVs. Due to the low numbers of archaea these were discarded from further analyses. Median read depth in the bacterial samples was 41000, with a range between 23000 and 118000. One bacterial sample failed to amplify DNA (6 reads total) and was discarded. Median read depth in the fungal samples was 23500, with a range between 5200 and 36000. After rarefaction median bacterial richness was 1195 (range between 818 and 2420 IQR 1064-1510) and median fungal richness was 264 (range between 147 and 470. IQR 225-296). The majority of the bacteria were within the proteobacteria and the acidobacteria phyla, while the majority of the fungi were in the sordariomycetes (supplementary figure 1).

### Impact of environmental properties

Microbial richness showed variable and limited responses to soil and plant environmental variables, while composition showed greater relationships (supplementary figure 2, Table 1). Fungal and bacterial richness showed differing responses to gradients in plant and soil properties, and were slightly negatively correlated with each other (Spearman rank correlation −0.05). Fungal richness increased with soil pH, and showed no relationship with plant richness or soil organic matter while bacterial richness decreased with soil pH and increased with both plant richness and soil organic matter (supplementary figure 2). Both bacterial and fungal richness had a small number of high diversity outliers, however these were not the same sites in both cases. The bacterial and fungal community compositions were analysed using NMDS ordination and each soil and plant variable fit to that ordination using a linear fit, which revealed that soil pH was the strongest predictor of bacterial and fungal community composition, with R^2^ of 0.85 and 0.65 for bacteria and fungi respectively (Figure 1, Table 1). Plant Ellenberg R (an index of plant responses to pH) and soil electrical conductivity (EC) were the next two most strongly associated variables for both bacteria and fungi (Table 1, see supplementary tables 1 and 2 for the full envfit results). For bacteria, whilst the first NMDS axis was largely explained by pH (Figure 1), soil structural components such as organic matter content, aggregate stability, and bulk density were orthogonal to the pH axis (Table 1). For fungi, pH also explained most of the variation along the first axis of the NMDS, whilst soil aggregate stability, clay content and legume species richness were orthogonal to the pH axis (Table 1). Soil organic matter and total carbon showed no significant relationship with fungal community composition, although they were both significantly associated with the second NMDS axis for the bacterial community. Some effect of pasture age is visible in the NMDS plot, especially once the gradient in soil pH is accounted for (Figure 1).

**Figure 1:**
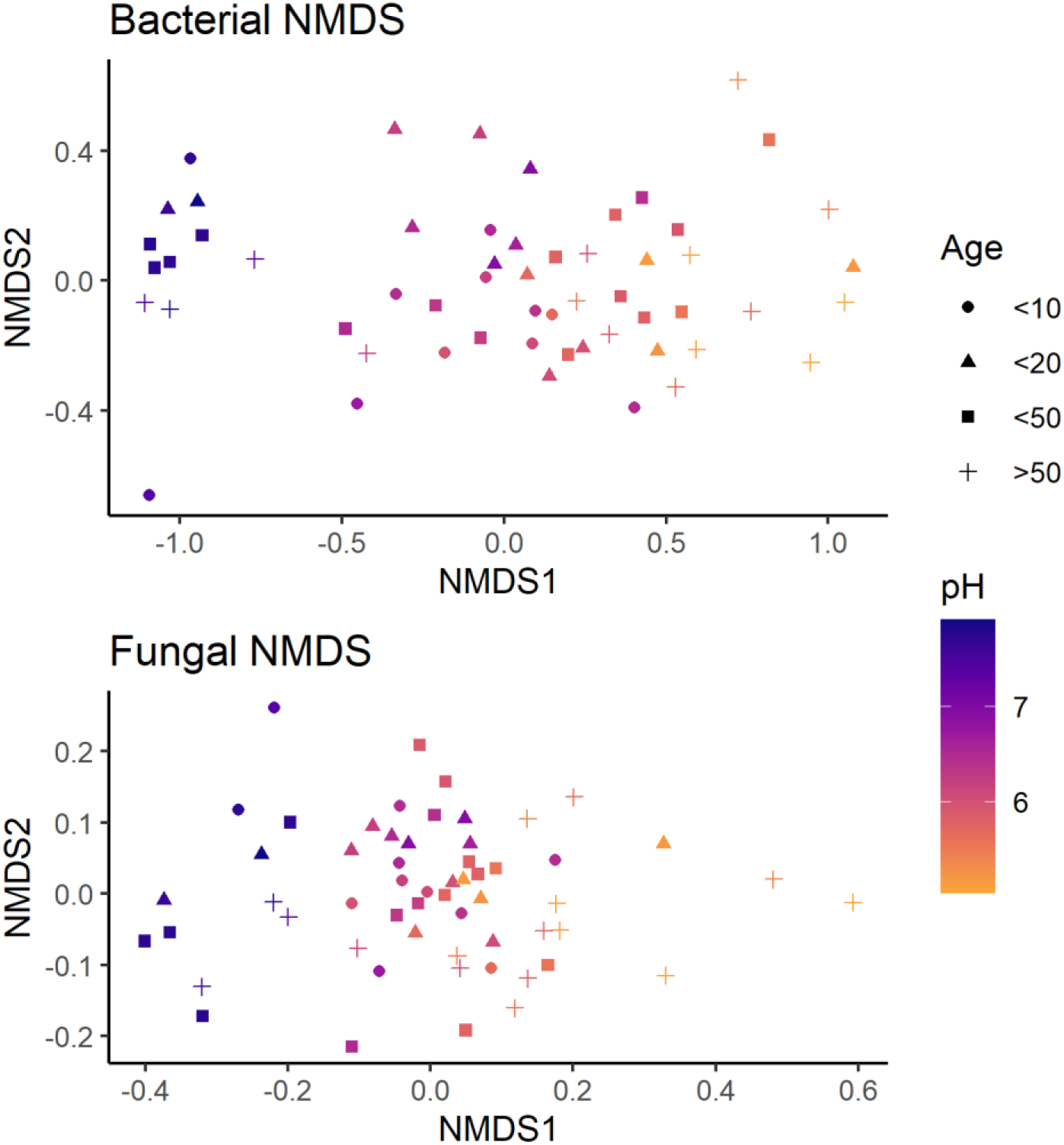
Ordinations of bacterial and fungal community compositions with the points coloured by pH and with shapes representative of the age of pasture. The bacterial NMDS stress was 0.11 and the fungal NMDS stress 0.17.

**Table 1:** Relationships between plant and soil parameters and the NMDS for bacteria and fungi as given by a linear regression upon the ordination scores, showing only variables that are correlated with either bacteria or fungi with R^2^ values greater than 0.15. For the full results see supplementary tables 1 and 2.

| Variable | Bacteria |  |  |  | Fungi |  |  |  |
| --- | --- | --- | --- | --- | --- | --- | --- | --- |
| | NMDS1 | NMDS2 | $R^2$ | p | NMDS1 | NMDS2 | $R^2$ | p |
| **pH (CaCl<sub>2</sub>)** | -0.965 | 0.262 | 0.849 | 0.001 | -0.979 | 0.206 | 0.651 | 0.001 |
| **pH (DIW)** | -0.954 | 0.300 | 0.809 | 0.001 | -0.955 | 0.296 | 0.643 | 0.001 |
| **Ellenberg R** | -0.675 | 0.738 | 0.450 | 0.001 | -0.995 | -0.103 | 0.480 | 0.001 |
| **Soil EC** | -0.847 | 0.532 | 0.371 | 0.001 | -0.981 | 0.194 | 0.315 | 0.001 |
| **Total N** | -0.264 | -0.965 | 0.364 | 0.001 | -0.869 | -0.495 | 0.059 | 0.237 |
| **LOI** | -0.195 | -0.981 | 0.353 | 0.001 | -0.827 | -0.562 | 0.022 | 0.588 |
| **Ellenberg F** | 0.221 | -0.975 | 0.276 | 0.002 | 0.504 | 0.864 | 0.102 | 0.065 |
| **Total C** | -0.348 | -0.938 | 0.274 | 0.002 | -0.999 | -0.034 | 0.055 | 0.273 |
| **Bulk density** | 0.076 | 0.997 | 0.194 | 0.009 | -0.068 | 0.998 | 0.026 | 0.531 |
| **Stability** | 0.326 | -0.946 | 0.152 | 0.016 | 0.314 | -0.949 | 0.228 | 0.003 |
| **Ellenberg N** | -0.725 | 0.689 | 0.114 | 0.055 | -0.618 | 0.786 | 0.301 | 0.001 |
| **Plant PCoA axis 1** | -0.507 | 0.862 | 0.111 | 0.055 | -0.499 | 0.866 | 0.267 | 0.001 |
| **Olsen P** | -0.972 | 0.237 | 0.074 | 0.177 | -0.487 | 0.873 | 0.206 | 0.008 |
| **Clay** | -0.248 | -0.969 | 0.096 | 0.088 | -0.166 | -0.986 | 0.159 | 0.018 |
| ***L. perenne* cover** | -0.603 | 0.798 | 0.047 | 0.311 | -0.392 | 0.920 | 0.197 | 0.006 |
| **Legume richness** | -0.135 | 0.991 | 0.032 | 0.468 | 0.046 | -0.999 | 0.198 | 0.006 |

Variation partitioning comparing the relative correlations of soil microbial composition with soil chemistry, soil structure, plant community and farm management showed that soil chemical properties (pH, total C, total N, Olsen P and conductivity) explained the largest amount of variation across both bacterial and fungal communities (Figure 2). The second largest proportion of variation was explained by the joint influence of plant community and soil chemistry, followed by soil chemistry and soil structure in both bacteria and fungi. Plant community composition also explained ~4% of variation directly for the fungal community while it showed limited influence on its own for bacteria. Soil structure (sand, silt, clay, aggregate stability) was also important for both bacterial and fungal communities, although to a lesser extent, explaining 1-2% of variation by itself and ~5% combined with soil chemistry. However, farm management properties as represented by the MCA ordination scores (pasture type, age, type of grazing, organic, winter grazing, fertiliser use, sheep presence) explained only a small amount of variation for fungi and bacteria and removing it from the variation partitioning increased the residual variation by only ~1 percentage point.

**Figure 2:**
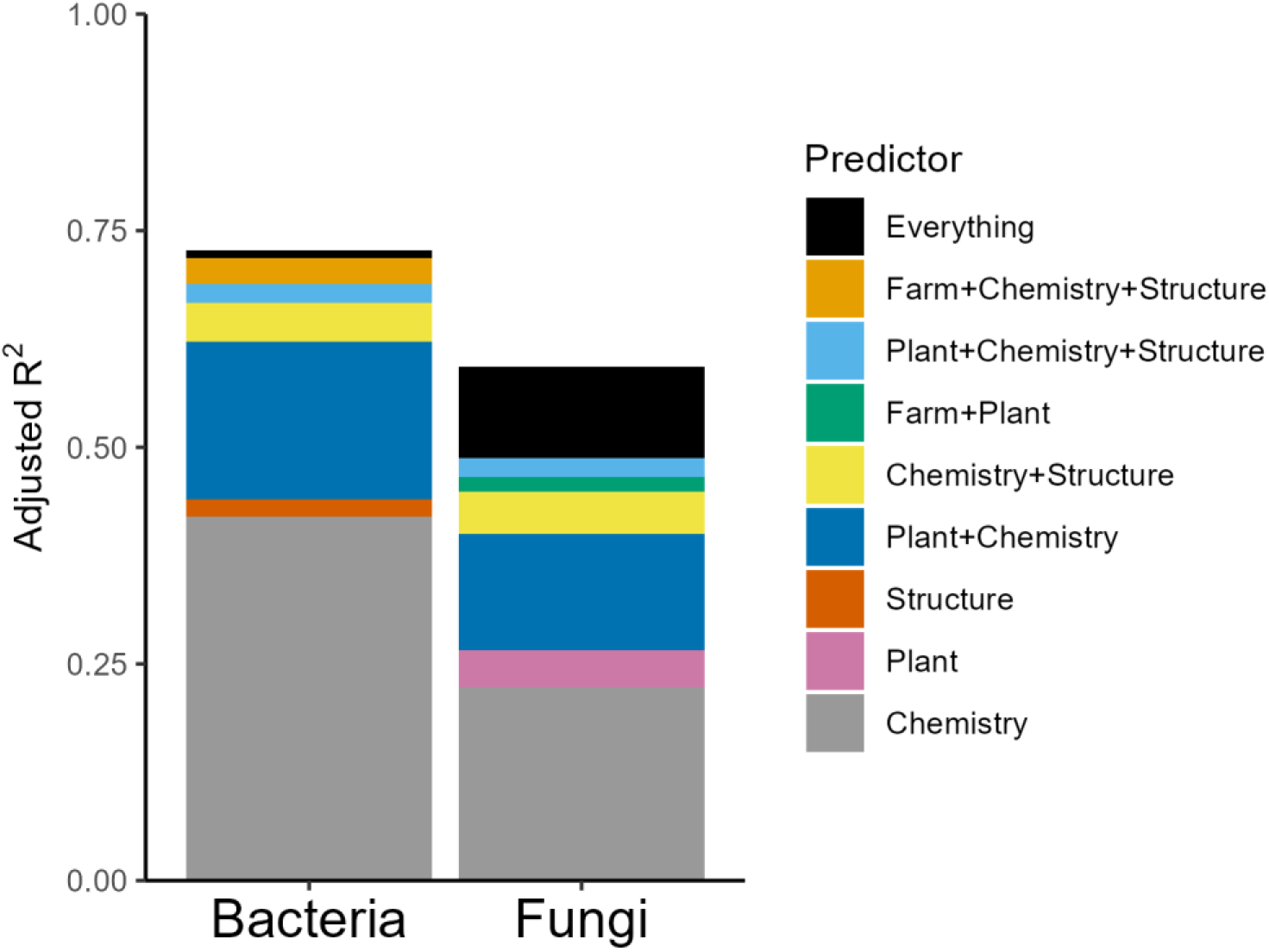
Proportion of variation in bacterial and fungal 4-dimensional NMDS scores explained by the different components. Chemistry and structure are the PCA scores of the soil chemical and structural components respectively. The plant component is represented by PCoA scores of the plant community and the farm component the MCA scores of key farm management variables including pasture age. Components that have multiple descriptors in the legend key are the joint variation explained by those named components together, and “Everything” is the variation explained by all of the components together.

Examination of the ordination scores for the soil chemical, soil structural, plant community and farm management data enables us to explore how these different components are represented within the variation partitioning presented in Figure 5. All five soil chemical properties measured were clearly represented in the ordination, with total C and total N showing a strong association with each other and pH, Olsen P and conductivity all being orthogonal to carbon in the first two axes and then each explaining independent variation in the third and fourth axes (Axes 1 to 4 explained 53%, 21%, 17% and 8% of variation respectively, Figure 3a, supplementary figure 3a). The soil structural parameters showed a clear separation primarily of the sand gradient (axis 1 explained 70% of variation), with aggregate stability being the major determinant of the orthogonal second axis (axis 2 explained 17% of variation, Figure 6b, supplementary figure 3b). The ordination of the plant communities showed that the major drivers of plant composition differences across the farms were differences in the covers of *Lolium perenne*, *Agrostis stolonifera*, *Trifolium repens* and *Holcus lanatus* (Relative eigenvalues of first four axes were 0.29, 0.11, 0.09 and 0.07 respectively, Figure 3c, supplementary figure 3c). The ordination of the farm management data achieved less explanatory power than the other three ordinations (first four axes explained 10%, 9%, 8% and 7% of variation respectively), as expected within MCA analysis, however the analysis did separate out organic farming on the first axis, older (>50 year old), often organically managed (if not certified), pastures on the second axis, and uses of lime and mineral fertilisers on the third and fourth axes respectively (Figure 3d, supplementary figure 3d). This indicates that the farm management effects included within the variation partitioning above are largely related to fertilisation, with a small inclusion of pasture age effects.

**Figure 3:**
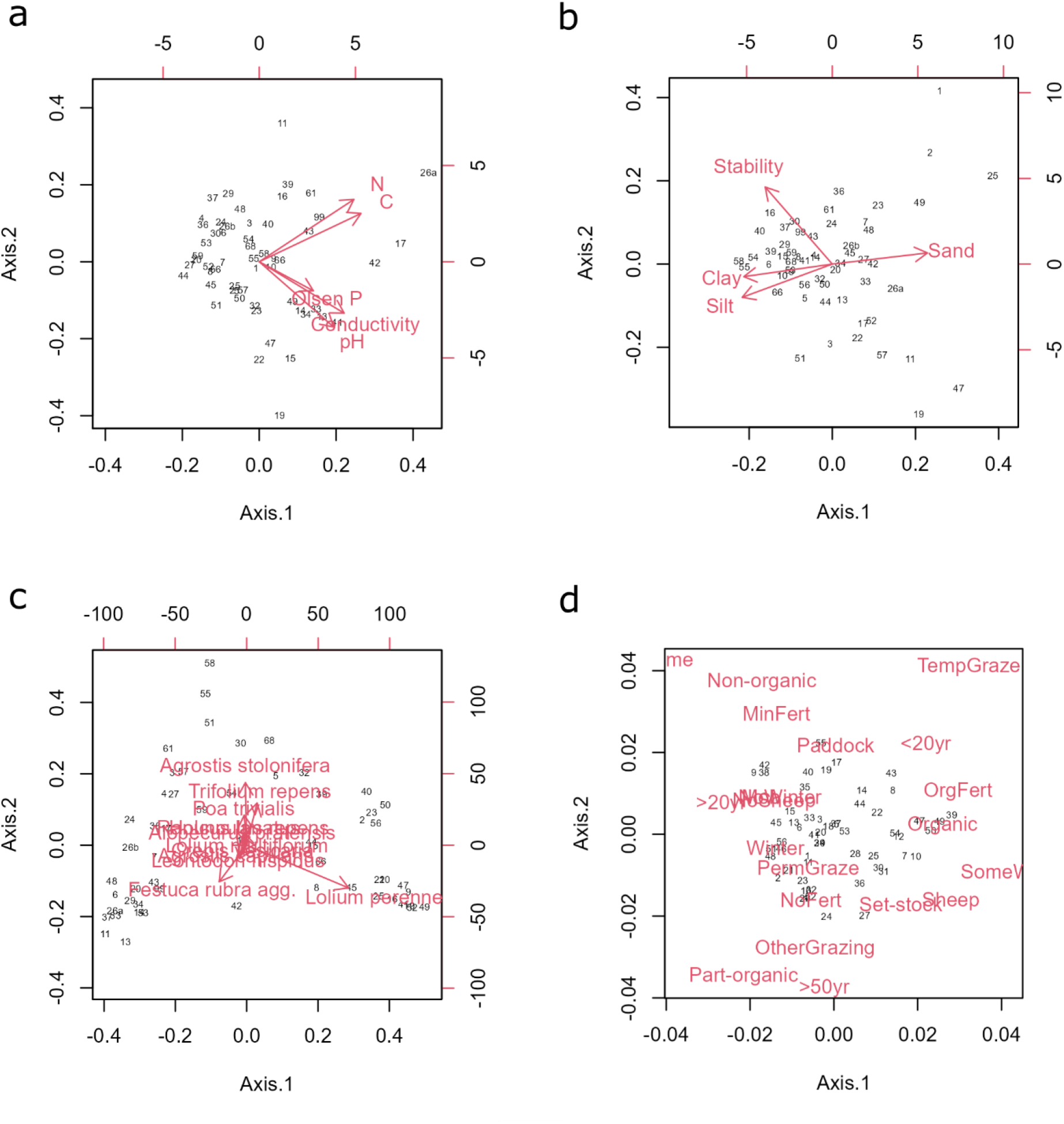
The first and second axes of the soil chemical PCA (a), soil structural (b), plant PCoA (c), and farm management MCA (d). Farms are represented by numbers, variable effects are represented by red arrows for a, b and c and centroids in d.

### Impact of land management

The direct relationship between farm management practices and soil physicochemical properties, vegetation composition or microbial communities varied depending on the environmental properties considered. Of the 57 fields sampled, 46 had been under the current land use (permanent pasture) for over 10 years with 16 of these farms having been under the current land use for over 50 years. There were also roughly equal proportions of mob grazing, set stocking and paddock/strip grazing (15, 18, and 16 fields respectively), and only a small number of fields receiving mineral fertiliser (10, compared to 19 receiving organic fertiliser). There were no significant trends with pasture age in soil organic matter, plant available phosphorus, cover of *Lolium perenne* or microbial richness (Figure 4). On average cover of *L*. *perenne*, Olsen P and fungal richness were lower in the oldest fields, however the 95% confidence interval of the monotonic age effect parameter crossed zero in all cases (*L*. *perenne* cover: −7.94 to 0.21, Olsen P: −3.86 to 0.65, fungal richness: −10.44 to 3.52). LOI and bacterial richness both had their 50% confidence interval cross zero. There were no trends in fungal trophic mode proportions over the different age categories (supplementary figure 4).

**Figure 4:**
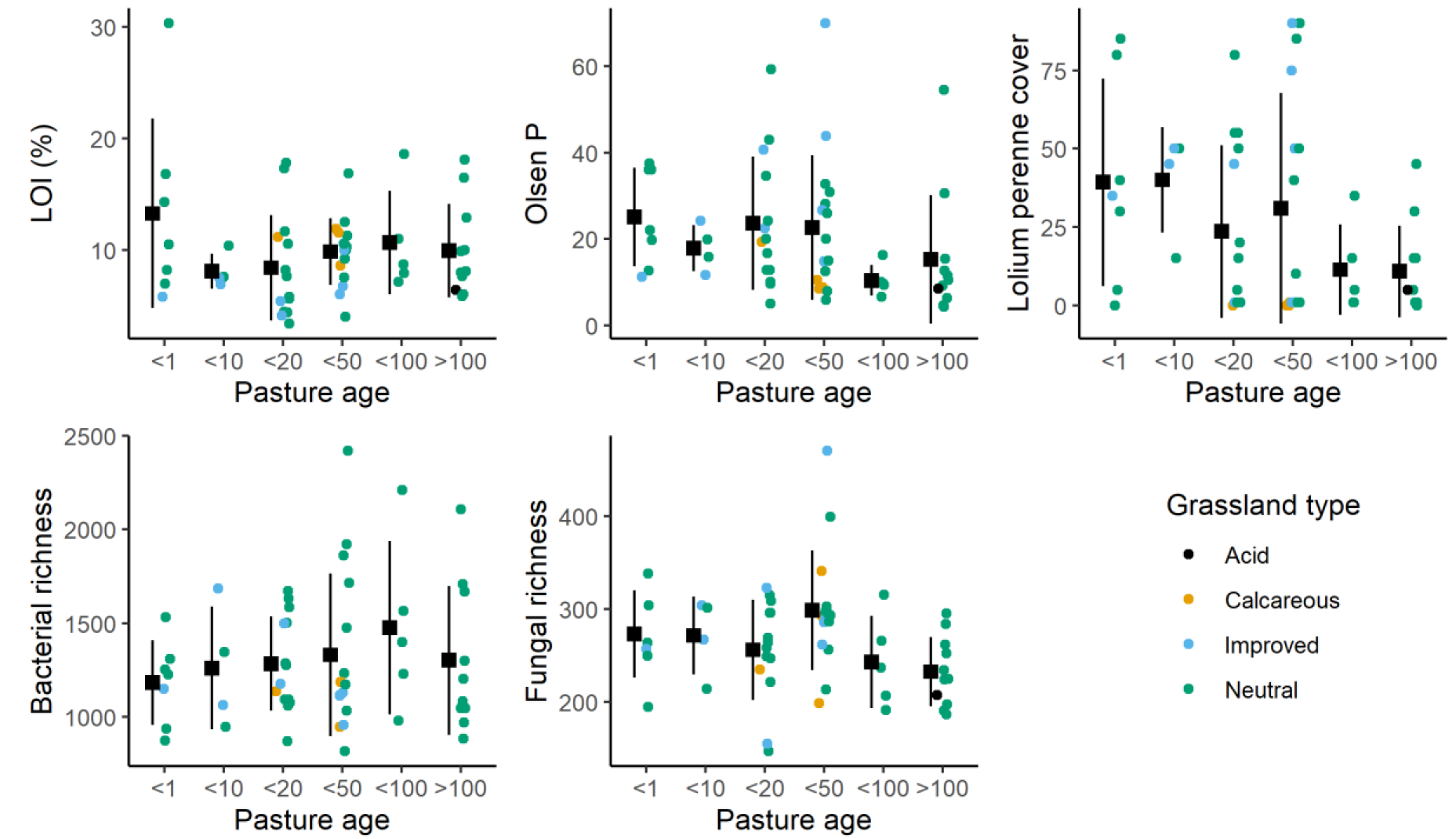
Differences in soil carbon, Olsen P, *Lolium perenne* cover, bacterial richness and fungal richness with pasture age and grassland type. Each field is represented by a single point, coloured by grassland type. The average, and standard deviation, of each field age category is represented by a black square and vertical lines.

Pasture age (<10, 10-20, 20-50 and >50 years) had no significant effect on bacterial community composition (dbRDA p = 0.10, Figure 5a). However, the fungal communities showed significant differences between field age categories (dbRDA p = 0.004, Figure 5b). Figure 5b shows a separation out of the youngest and very oldest pastures, the latter perhaps being partly driven by higher variance in the oldest category of farms (betadisper p = 0.04). There is more separation by age apparent in Figure 5 compared to Figure 1 due to dbRDA being a method of constrained ordination, versus NMDS being unconstrained ordination, and therefore representing changes in microbial composition that are not occurring in the two primary axes of overall variation. There was no significant impact of grazing type (mob, set stocking or paddock/strip) on either bacterial or fungal communities (dbRDA, p = 0.43 and p = 0.16 respectively, supplementary figure 5). Fertilisation regime (mineral, organic or none) had no significant effect on bacterial composition but did show differences in the fungal composition (p = 0.10 and p = 0.035 respectively, supplementary figure 6). These results also reflect the higher variance in community composition across the no fertiliser fields, which comprised 57% of the fields sampled (betadisper p < 0.001).

**Figure 5:**
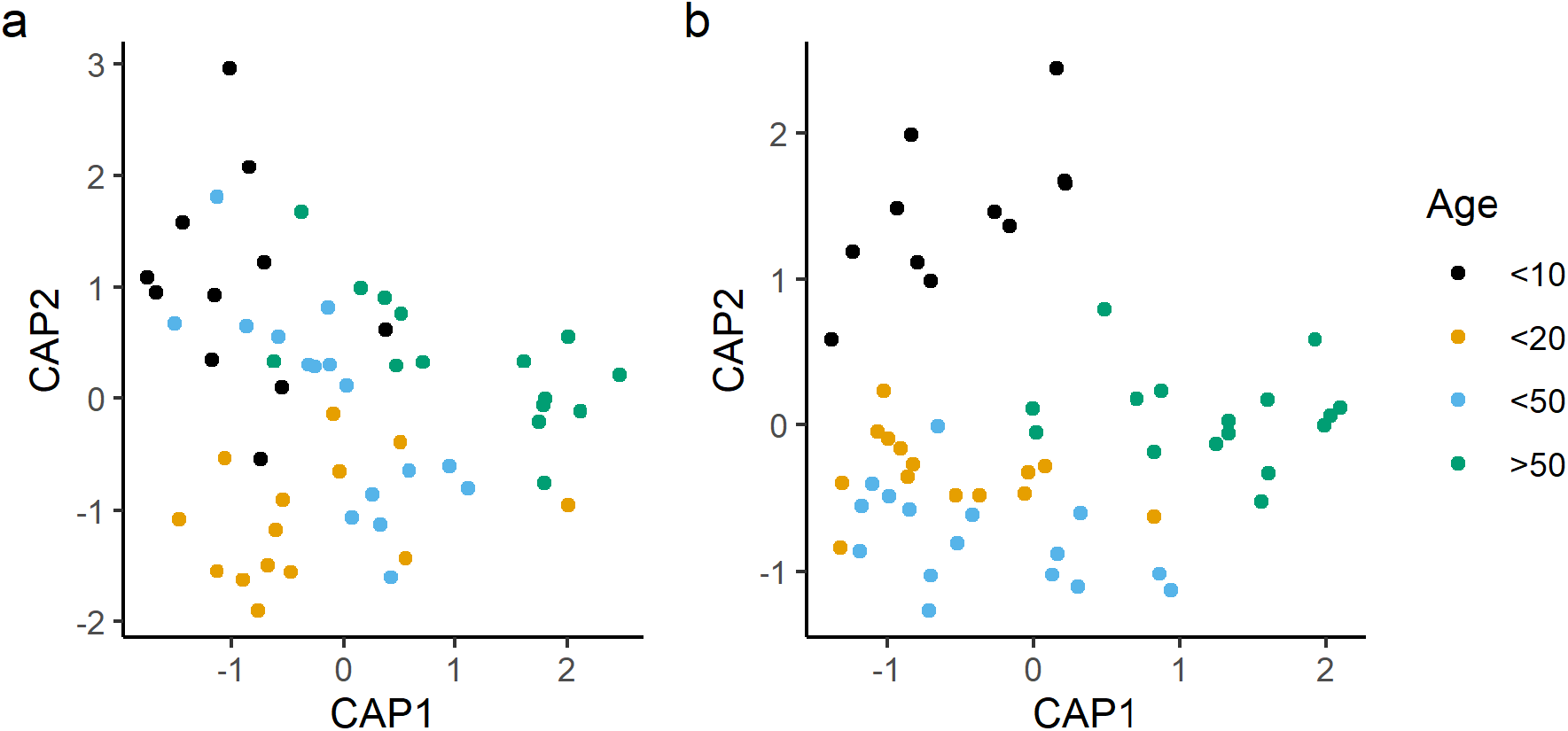
The first two axes of the dbRDA based on age of pasture for bacteria (a) and fungi (b)

### Indicator taxa

Indicator taxa analysis identified hundreds of bacterial and fungal taxa as potential indicators of pasture age once differences in soil pH were accounted for (868 bacterial taxa and 450 fungal taxa, corresponding to 2.8% and 6.7% of all taxa respectively). Both bacteria and fungi had more taxa that responded to the pasture age cut-off of >50 years old than that of >20 years old. Some of these taxa also responded to changes in carbon, Olsen P or *Lolium perenne* cover once soil pH had been taken into account (Figure 6). However, these trends were not always consistent, for example there were hundreds of taxa that were indicative of both younger fields and differences in carbon concentration but some of these increased with increasing carbon and some decreased (~40% fungi decreased and ~20% bacteria decreased). The only bacterial taxon which was higher in both older farm categories and also higher in organic carbon soils was a member of the Micromonosporaceae family, and also decreased in response to *L*. *perenne* cover and Olsen P content. The taxonomic breakdown of the indicators was broadly similar to that of all the taxa, with read counts over 10 for both bacterial and fungal classes, however the Agaricomycetes made up 23% of the PFLA quality indicators compared to 16% of the PFLA data overall, and 17% of the PFLA data with read count above 10 (supplementary figure 7). Overall the proportion of Agaricomycetes was positively associated with LOI, unlike the other four most common fungal classes and the Glomeromycetes (Figure 7).

**Figure 6:**
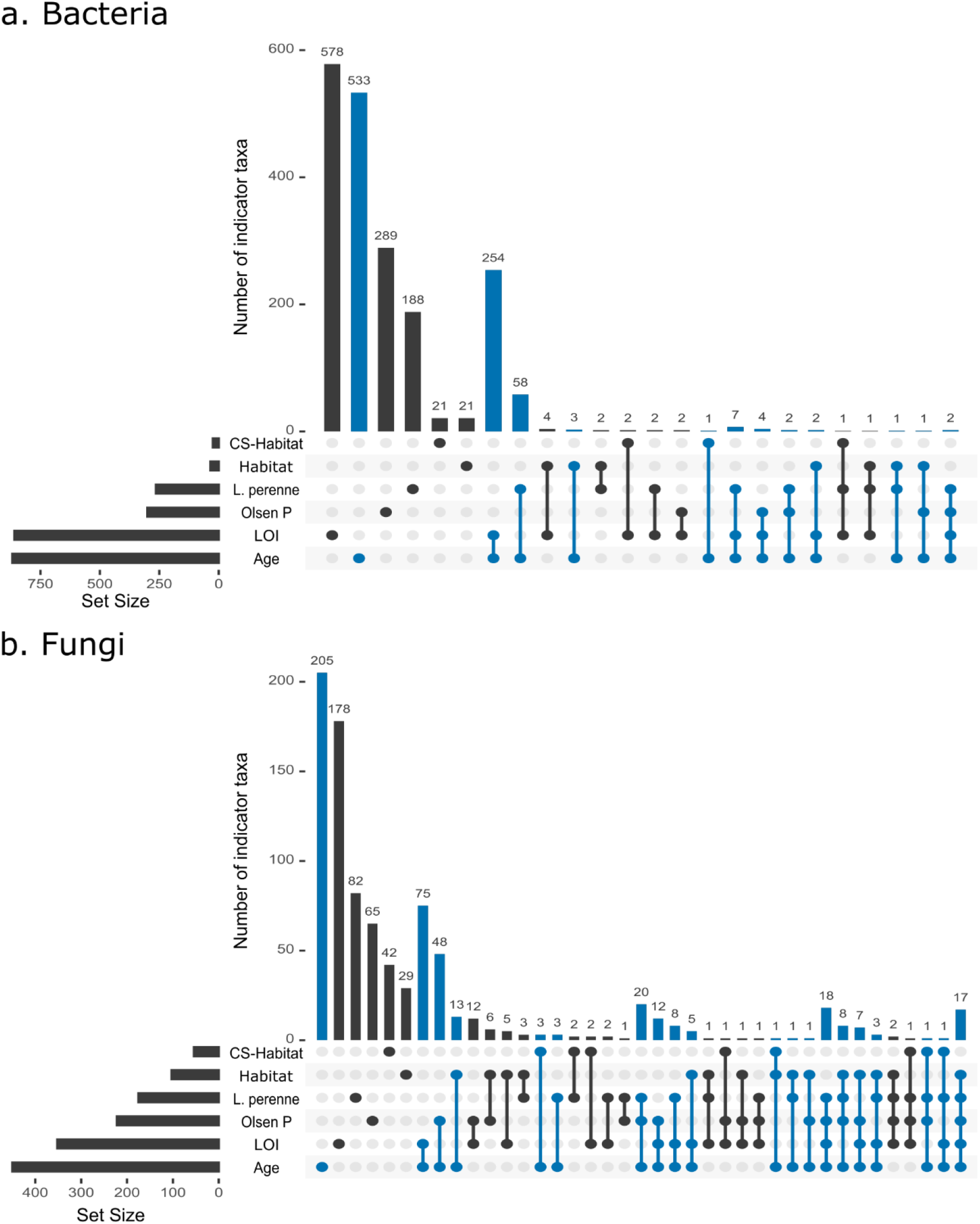
The counts of the different overlaps between the indicators of habitat, *L*. *perenne* cover, Olsen P, LOI and pasture age once the pH gradient is accounted for in bacterial (top) and fungal (bottom) communities represented by an upset plot. The total number of taxa that significantly respond to each soil or farm quality indicator is given by the horizontal bars to the left of the plot. Each vertical line within the plot represents a certain overlap of indicator taxa groups, the overlap identity is given by the conjoined dots while the total number of taxa within that overlap is given by the length of the vertical bar and the number above the bar. For example, the leftmost vertical bar in panel b (coloured b) represents the 205 fungal taxa that respond solely to pasture age (indicated by the single blue dot within the age row in the below matrix) while the rightmost vertical bar represents the 17 fungal taxa that respond to habitat, *L*. *perenne*, Olsen P, LOI and pasture age (indicated by the conjoined blue dots bridging the five rows of the below matrix). Both of these vertical bars are included in the total indicator for age count in the horizontal bar to the right. Indicator overlaps that respond to age are coloured blue, all others are coloured black.

**Figure 7:**
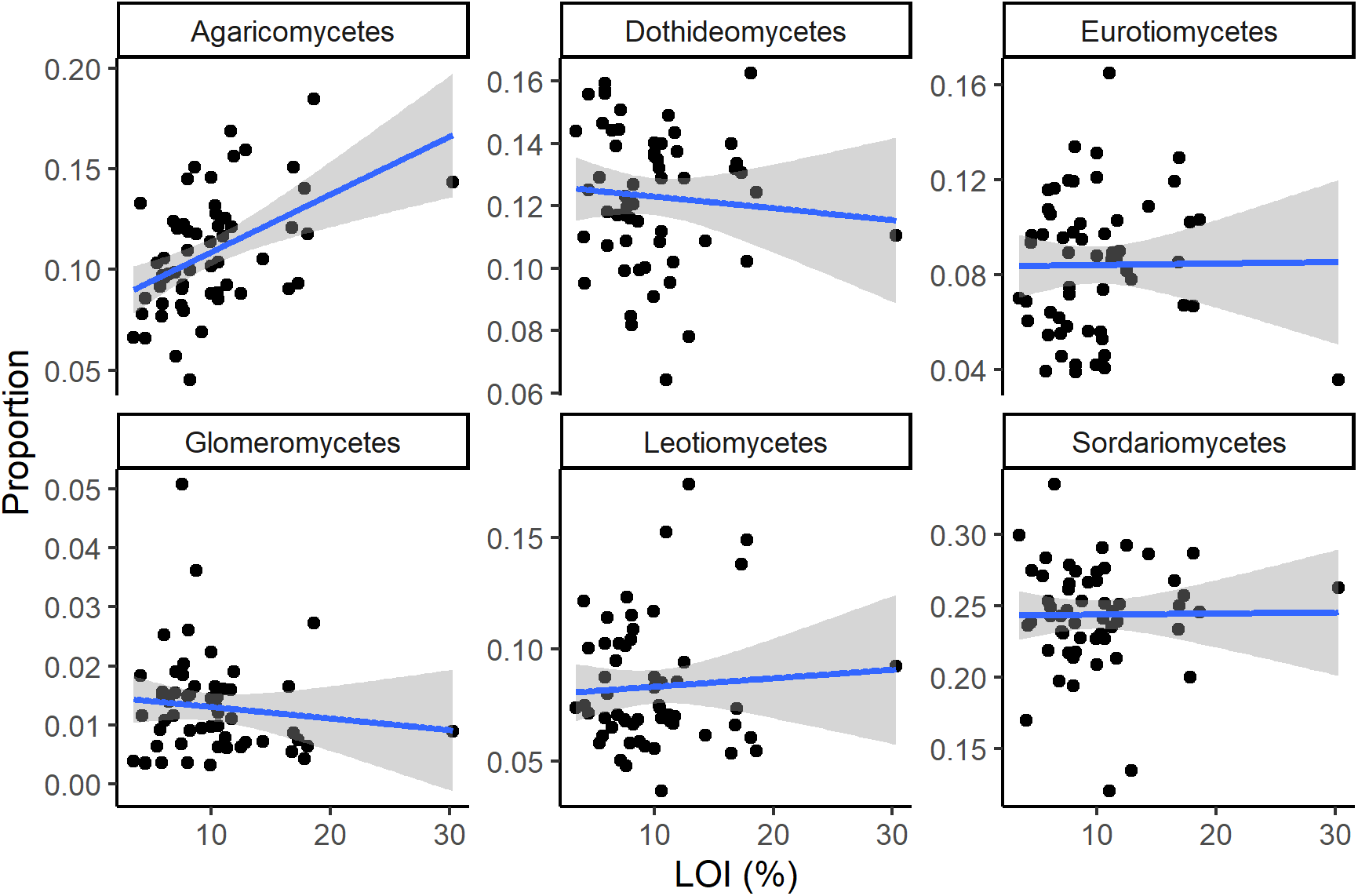
The differences in fungal class proportion across the LOI gradient. The agaricomycetes were positively associated with LOI (R^2^ = 0.22, p < 0.001 with and without the high LOI outlier), while all other relationships were non-significant (p values of 0.55, 0.94, 0.43, 0.64 and 0.94 for each class in alphabetical/panel order).

### Comparison to grassland soils in the wider GB landscape

In order to assess the generality of the indicators of soil health identified above the PFLA indicators were compared to indicators of grassland type within the Countryside Survey, of which there were 23 bacterial and 127 fungal taxa that showed differences between Neutral and Improved grassland once pH was accounted for. Of these, 12 bacterial taxa and 76 fungal taxa were more abundant in Neutral grassland than in Improved. A small number of these taxa also showed responses to either field age or other properties potentially indicative of field quality within the PFLA data (Figure 6). Within the bacterial data there were only 4 taxa that showed a response to both CS grassland type and a PFLA soil property, this included two Acidimicrobiales which showed higher prevalence in farms over 50 and soils with higher organic carbon and matched at 100 and 99% sequence similarity respectively to a CS taxa that showed higher prevalence in Neutral grassland. The other two taxa were more prevalent in Improved grassland in CS, one a member of the SMB53 genus was less common at higher levels of organic carbon and the other a member of the MBNT15 order was more common where there was high organic carbon and Olsen P.

Within the fungal data there were many more common indicators of grassland type and soil quality, however these often showed contradictory responses to the differing soil quality indicators. Both *Cuphophyllus borealis* a facultative saprotroph/Symbiotroph and *Gibellulopsis piscis* a probable facultative plant pathogen preferred Improved to Neutral grassland in CS and were less commonly found in older farms (>50 years) in PFLA. However, *C. borealis* was also less common at high Olsen P and *L*. *perenne* cover in the PFLA farms. *Leohumicola minima* and *Fusarium culmoram* both showed preference for Improved grassland in CS and increasing *L*. *perenne* cover in PFLA. *Thelebolus spongiae* (probable saprotroph/Symbiotroph) showed a preference for Improved grassland in CS and lower organic carbon in PFLA; while a member of the Clavaria genus showed a positive response to Neutral grassland in CS and higher organic carbon in PFLA. All other taxa that were indicators of both grassland types in CS and a field quality property in PFLA were only identified to coarser taxonomic levels. The five taxa that showed higher presence in Neutral grassland compared to Improved grassland plots in CS and also responded to a PFLA property (increasing with age or carbon) were all members of the Agaricomycetes, with the 4 identified to order level being Agaricales (this includes the Clavaria identified above). All others were not identified to a more detailed taxonomic level. However, it should also be noted that Agaricales is a common order, and in fact 13 Agaricale taxa showed higher prevalence in Improved grassland in CS while 28 showed higher prevalence in Neutral grassland. None of the 13 Agaricale taxa that were more abundant in Improved grassland also responded to a property within the PFLA data.

## Discussion

Across the farms within our analysis we found pasture age since plough impacted fungal community structure, as well as specific bacterial and fungal taxa, while fertilisation and grazing regime showed lower impacts. Fungal community composition, and to a lesser extent richness, showed trends over the pasture age gradient sampled. The stronger relationship between pasture age and microbial composition compared to microbial diversity is consistent with previous findings showing greater changes in microbiome composition versus diversity in response to soil management (Neal et al., 2021). There were also hundreds of taxa identified as being more abundant in either younger or older pastures. In particular there were greater differences in microbial taxa in pastures that had been established for over 50 years compared to younger pastures, demonstrating the importance of long-term farm management in determining microbial community structure. Pasture age influences plant rooting, community structure, soil structure and chemical properties (Faville et al., 2020; Löfgren et al., 2020; Tozer et al., 2016). In longer established pastures plant/soil interactions are better established, and we also found plant and soil chemistry interactions to be important for both bacterial and fungal community composition. The age gradient across our fields represents a variety of potential soil and plant changes, which will be modulated by farming practices as well as local environmental conditions.

Variability in grassland management practices, such as fertiliser applications, across the fields may obscure the changes we’d expect to see over the pasture age gradient, however we found only limited evidence for effects of management decisions such as grazing intensity and fertilisation inputs. This perhaps reflected the practices supported by the PFLA and the fact that only a minority of plots were classified as Improved grassland (9 out of 56), on which you would expect practices such as liming and fertilisation to influence soil properties and plant communities (Eze et al., 2018; Goulding, 2016). Our variation partitioning results do indicate that any impact of liming or fertilisation upon bacteria and fungi are likely fully mediated by changes in the soil physicochemical properties. Therefore, while we have not found strong evidence for a direct link between fertilisation and liming and microbial structure this does not necessarily mean that changes in fertilisation and liming are having no effect, as these management practices influence soil chemistry which we have found to be strongly associated with microbial community structure. It is likely that the impacts of fertilisation and liming upon soil chemistry and thus microbial community are influenced by the underlying soil type and climatic conditions at each farm (Goulding, 2016). Previous analysis has found limited effect of rotational grazing or differences in stocking rates upon soil properties, which was consistent with our lack of evidence for an impact of grazing regime upon microbial communities (Pyle et al., 2019).

Pasture age had limited effects upon plant and soil properties within our dataset, likely due to the range in environmental conditions covered by our survey. Previous studies have found declines in soil available phosphorus with increasing pasture age (Asner et al., 2004; Löfgren et al., 2020). While the trend was non-significant we did find lower Olsen P in older fields, likely due to the length of time since fertilisation. The lack of consistent trend in soil organic matter with pasture age may be related to the high variability in soil types and farm location, which are known to have big impacts on soil carbon (Hewins et al., 2018). While pasture has been found to have higher soil carbon than arable fields (Lin et al., 2020), previous examinations of soil carbon stocks over a range of pastures of different ages have found no significant effect of pasture age upon soil carbon due to variability in soil parent material and management practices, in line with our results (Derner & Schuman, 2007; Orgill et al., 2015). Other studies which have shown a decrease in soil carbon with pasture age are less comparable as the previous land cover was woodland, and the woodland to pasture transition is markedly different from the within-pasture types and arable to pasture transition, generally associated with increases in soil carbon which are common to our sites (Asner et al., 2004; Lin et al., 2020).

Soil chemical properties proved to be the key determinant of bacterial and fungal richness and composition within our data, although some soil structural and plant metrics also showed associations with microbial composition. Consistent with many previous studies of soil bacterial and fungal communities, soil pH was the variable most associated with community composition (George et al., 2019; Griffiths et al., 2011). Surprisingly, while soil carbon showed some association with the bacterial communities there was no relationship between soil carbon and fungi. We also found that soil electrical conductivity was strongly associated to both bacterial and fungal communities. This may be related to the fact that soil solution composition has a direct effect in the attractive/repulsive forces at the clay scale causing the clay particles to come together into microaggregates, the higher the electrical conductivity the more stable the aggregates will be, providing a better substrate for the microfauna to thrive (Lebron & Suarez, 1992; Shainberg & Letey, 1984). Soil structure showed some influence on the bacterial and fungal communities, in particular aggregate stability showed associations with the 2^nd^ axis of the ordination for both bacteria and fungi indicating that the soil structural influence on microbial compositions is orthogonal to the pH influence. Aggregate stability is known to be related to microbial communities and their activity, and while we have no detailed pore structural information it is likely that this is a strong determinant of microbial community structure and activity in our soils (Tecon & Or, 2017). It is also plausible that bacterial and fungal communities are influencing soil structure and soil chemistry, through carrying out a variety of functional processes and resulting in soil-microbe feedback mechanisms (Frac et al., 2018; Neal et al., 2021; Totsche et al., 2010).

While we found only some evidence for direct associations between the plant community and the microbial community we found that interactions between the plant community and soil chemistry were very important in determining microbial community structure. These interactions may reflect the difficulty in separating out the inter-correlated responses of soil properties and plant communities to environmental and management factors. For example, Löfgren et al. (2020) found that changes in plant communities over a >200 year pasture age gradient were often associated with changes in soil chemistry, particularly phosphorus and to a lesser extent nitrogen. It has also been found that plant community properties are more representative of historical, rather than current, soil conditions, particularly in metrics such as the Ellenberg scores (Wamelink et al., 2002). Therefore, the plant-soil interactions we have found to be important in determining bacterial and fungal community structure could be better representing an integrated long-term soil condition measurement than true plant-soil interactions, e.g. Ellenberg N could be better associated with the average soil fertility over the past few years rather than the current soil fertility.

We found that fungi were more responsive than bacteria to various factors including: management, plants and soil chemistry. Several of these factors interacted with each other, as identified within our variation partitioning analysis (Figure 5) leading to the need to condition analysis upon the differences in soil pH in order to identify microbial taxa that respond to soil quality indicators across the wide range of farms. This is consistent with our understanding that the impact of farm management decisions is context-dependent (Hannula et al., 2021; Kibblewhite et al., 2008). Soil pH was more associated with bacterial composition than fungal composition, consistent with previous studies of microbial composition in temperate habitats (George et al., 2019). Broad-scale bacterial community composition was better explained within the variation partitioning than broad-scale fungal community composition. Bray-Curtis distance responds more strongly to the dominant members of the community (McKnight et al., 2019), and it is possible that strong effects of soil chemistry and pH upon soil bacterial composition are driven by a relatively small number of pH-responsive dominant bacterial taxa. Notably, there were proportionally many more fungi than bacteria that responded to multiple drivers within the farms in the indicator analysis (Figure 7). There were also several unexpected and inconsistent responses of indicators to the different pasture quality indicators, where a single taxa responded in opposite directions to two indicators, potentially relating to relatively narrow regions of optimal niche space for the microbial taxa and suggesting routes for further investigation. Hence, changes in the soil fungal community and key fungal taxa may prove a better integrator of soil condition in pasture farms than changes in the bacterial community.

Within the indicator analysis the Agaricomycetes, and particularly the Agaricales, emerged as a particularly responsive group to changes in key pasture health-related properties such as soil organic matter. The agaricomycetes are ecologically diverse, including a variety of mushroom-forming fungi that obtain nutrients from saprotrophy, symbiotrophy and the occasional pathotrophy, including many ectomycorrhizal taxa (Sánchez-García et al., 2020; Tedersoo et al., 2010). Ectomycorrhizal fungi are important to various aspects of plant and soil health, as they can enhance nutrient uptake, protect plants against pathogens and pollutants, and also have various implications for soil carbon cycling (Kumar & Atri, 2018; Tedersoo & Bahram, 2019). While the various agaricomycetes are responding differently to different properties it may be possible to identify a small number of different taxa as key indicators of soil condition. As agaricomycetes are mushroom forming, this offers the opportunity for in-field mushroom surveys to evaluate soil condition, previously suggested as a method for evaluating forest health (Egli, 2011).

The agricultural sector in the UK and elsewhere is currently experiencing drastic changes related to both changes in consumer habitats and subsidy structure. Both cattle and sheep numbers have decreased in the UK in the past few decades (Defra, 2021), and these farms have been judged to be more at risk from changes to agricultural subsidies than arable farms (Arnott et al., 2019). Farmers show large variation in both their exposure to innovative farming practices and their willingness to change (Arnott et al., 2021). The farms we have surveyed within this work are part of the Pasture for Life Association which promotes innovative grazing techniques and brings together farmers that engage in practices that promote ecosystem health (Vetter, 2020). They represent a range of sustainable farm management practices across a wide range of environmental conditions, and our results show how the impact of management decisions upon soil bacteria and fungi are dependent upon, and mediated by, the soil chemistry. This highlights the importance of considering local conditions when monitoring and evaluating the impact of farm management on soil health. Our results also indicate that we have to consider the longevity of sustainable farming practices, and provide long-term stable support to maintain sustainable farming practices over the timescales required to support soil health.

## Acknowledgements

We thank the PFLA and all the farmers involved in the survey described for working with us on this project. We also thank land-owners, surveyors and project staff involved in Countryside Survey.

## Supplementary materials

**Figure 1:**
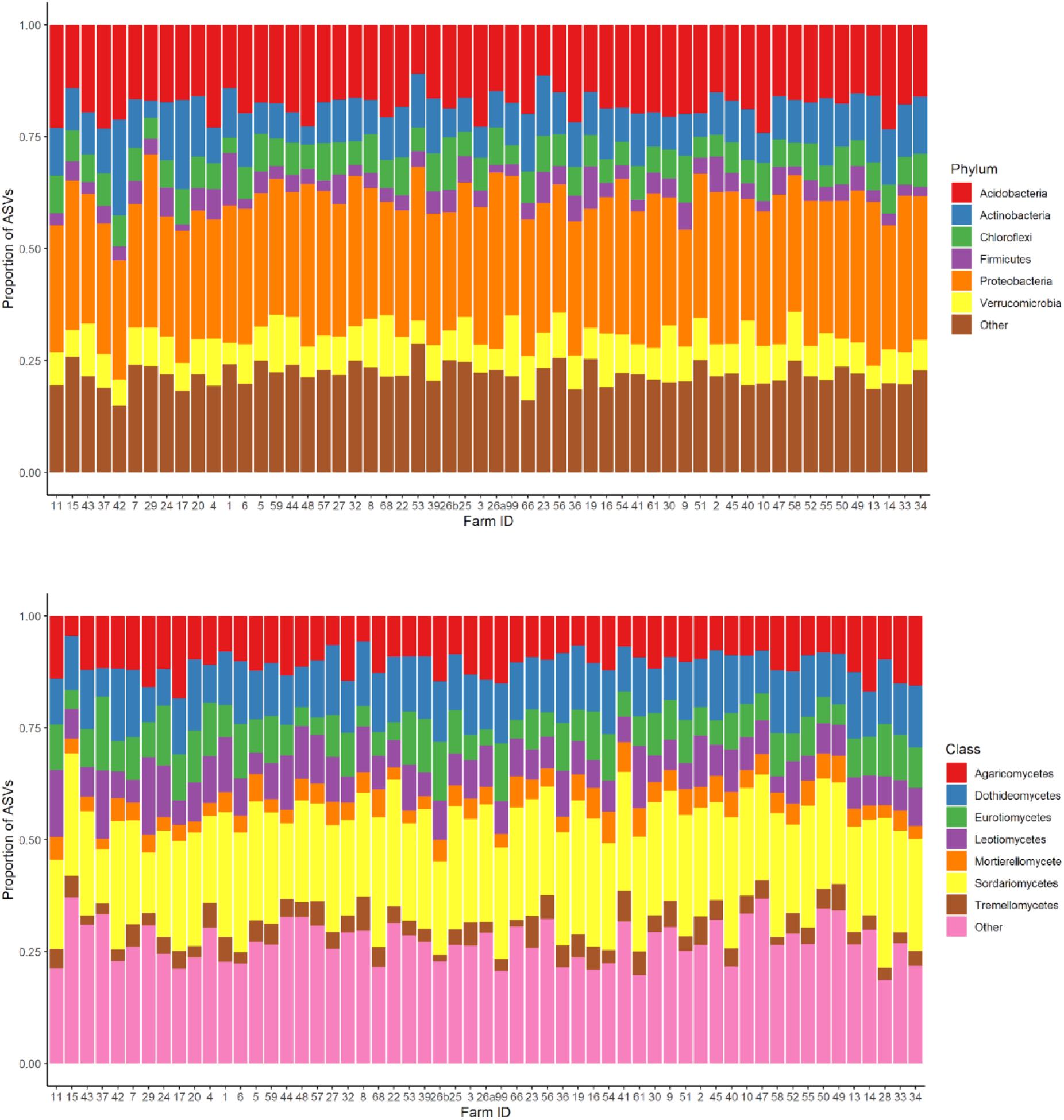
Bacterial phyla (top) and fungal class distributions across the farms, with farms ordered along the x axis from high to low plant community species richness/other quality indicators

**Figure 2:**
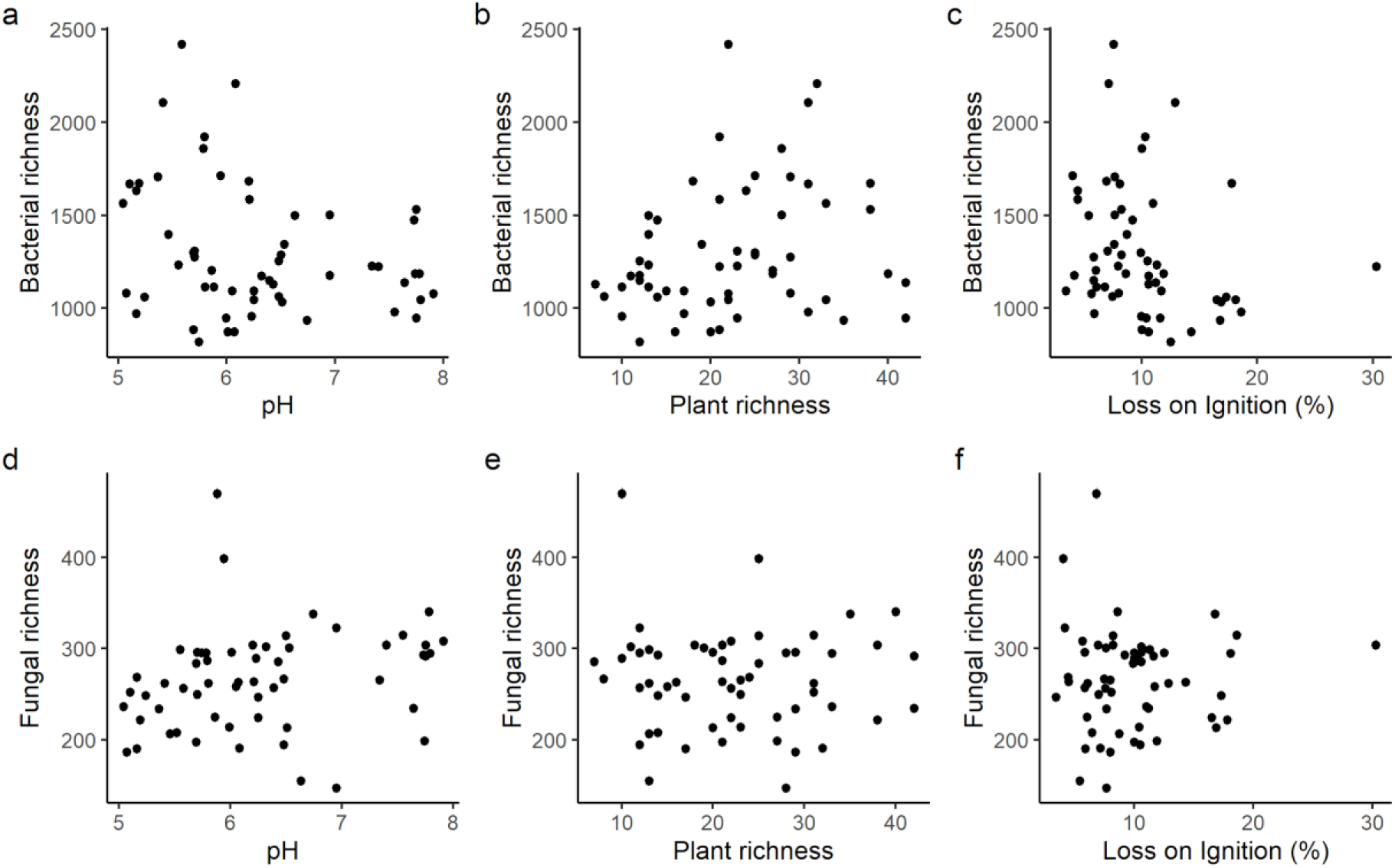
Bacterial and fungal richness against soil pH (a,d), plant richness (b,e) and organic carbon (c,f). Spearman rank correlations were −0.25, 0.35, −0.32, 0.37, −0.07 and 0.03 for a to f respectively.

**Figure 3:**
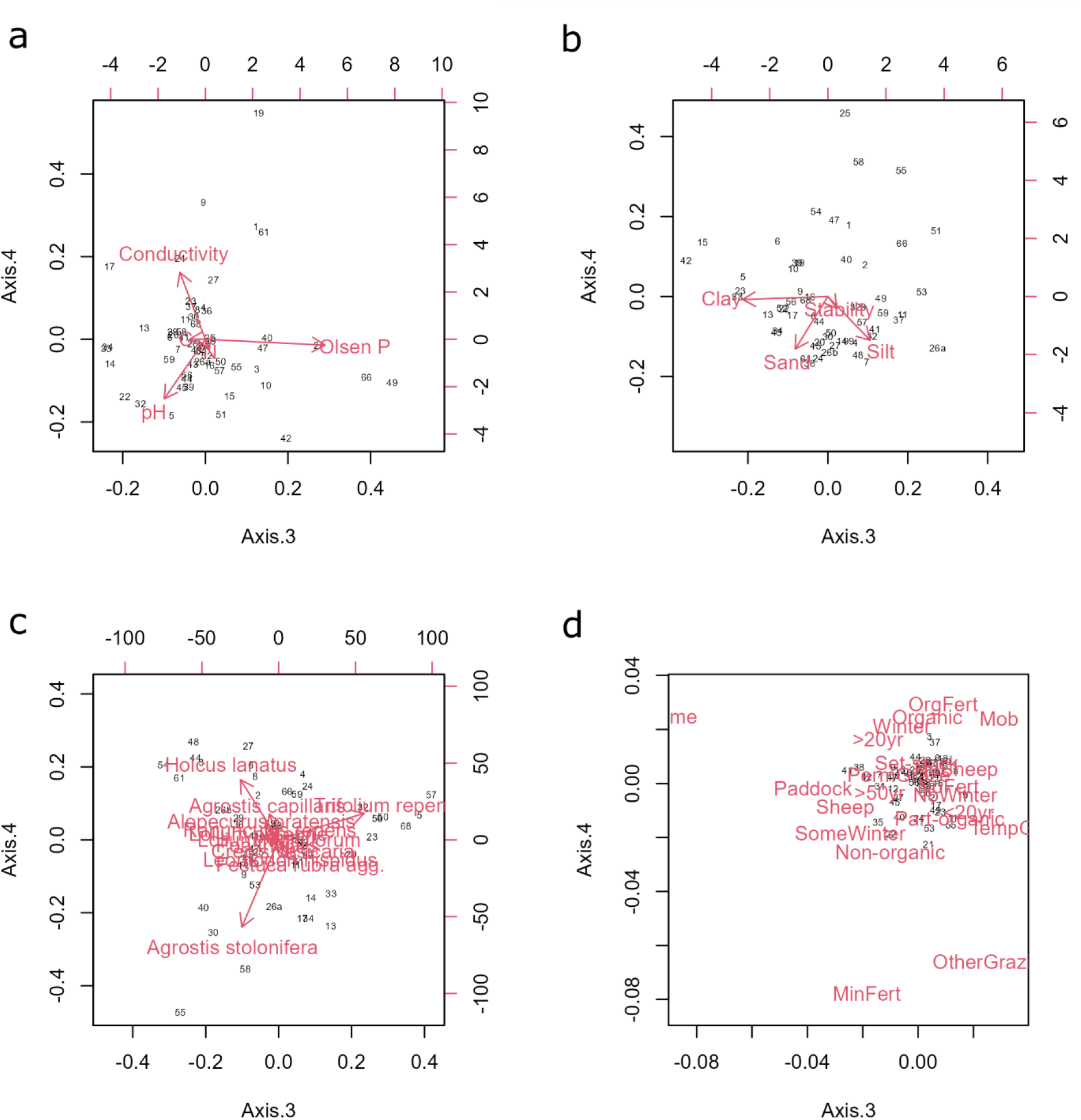
The third and fourth axes of the soil chemical PCA (a), soil structural (b), plant PCoA (c), and farm management MCA (d). Farms are represented by numbers, variable effects are represented by red arrows for a, b and c and centroids in d.

**Figure 4:**
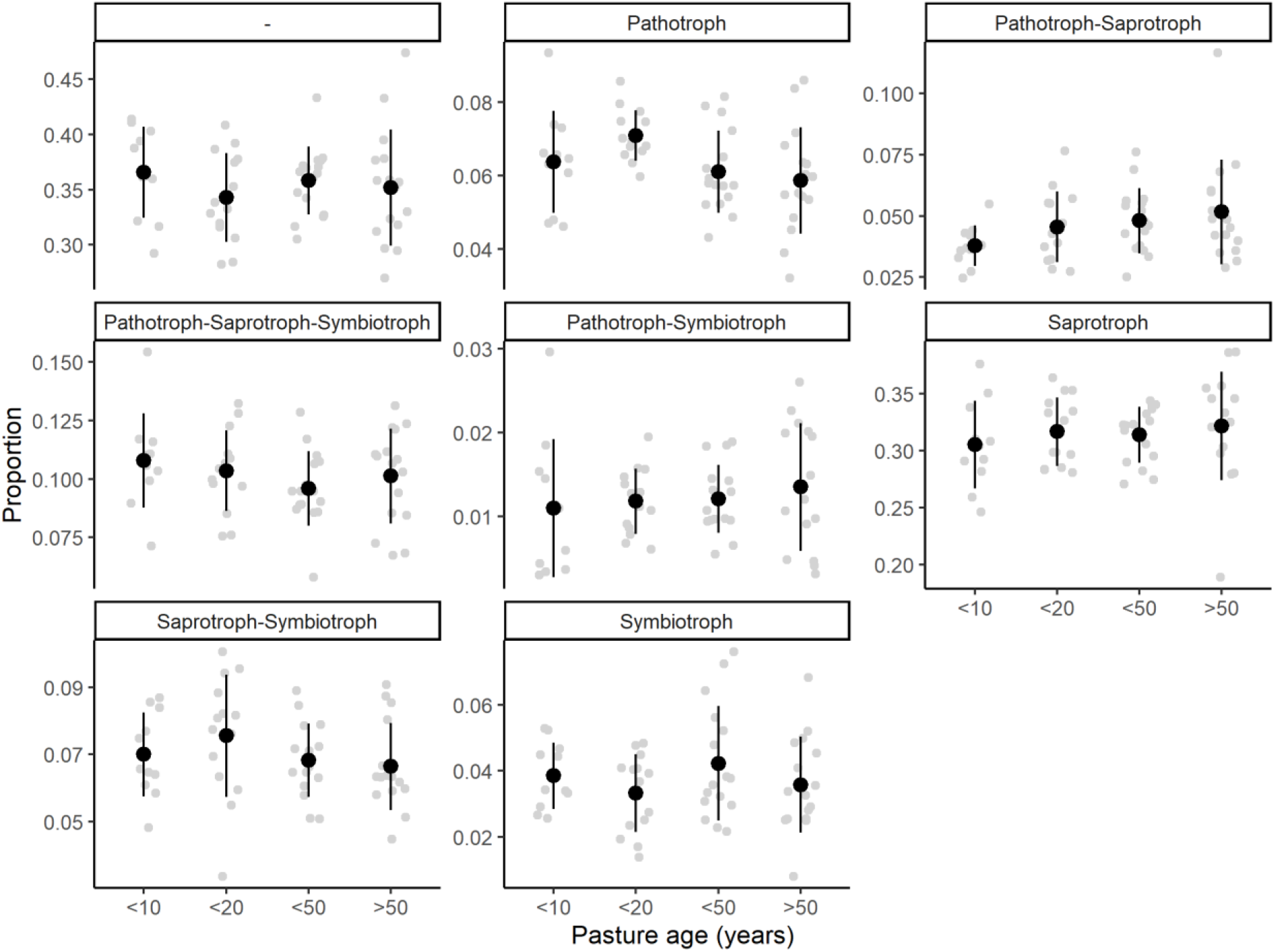
Fungal trophic mode proportion by pasture age. The mean and standard deviation are shown by black dots and vertical lines, with each field being represented by a grey dot randomly jittered on the x axis behind.

**Figure 5:**
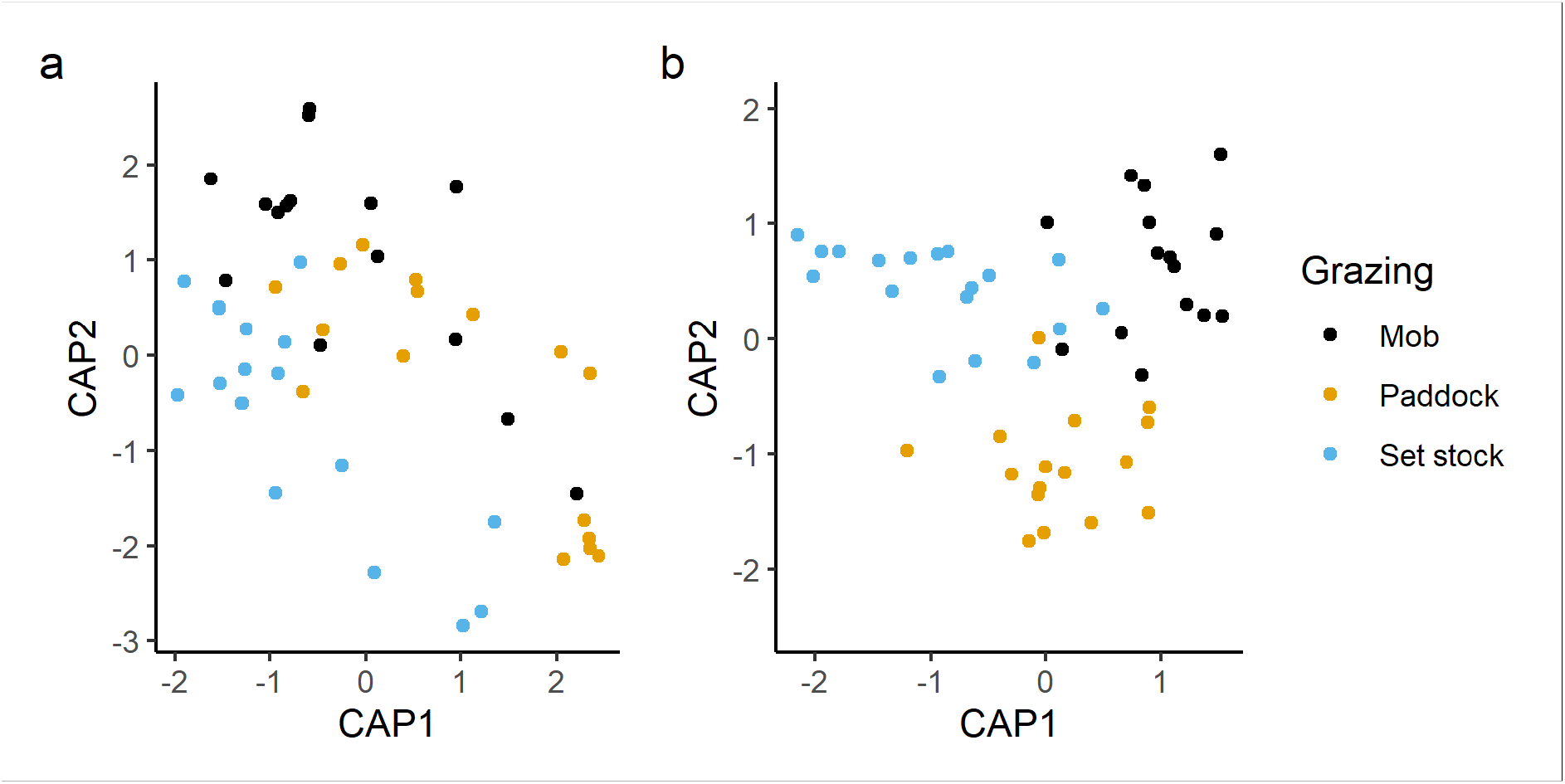
The first two axes of the dbRDA of the bacterial (a) and fungal (b) communities regressed against grazing regime. Results were non-significant with F_2,46_ = 0.990 and p = 0.427 and F_2,46_ = 1.105 and p = 0.166 for fungi.

**Figure 6:**
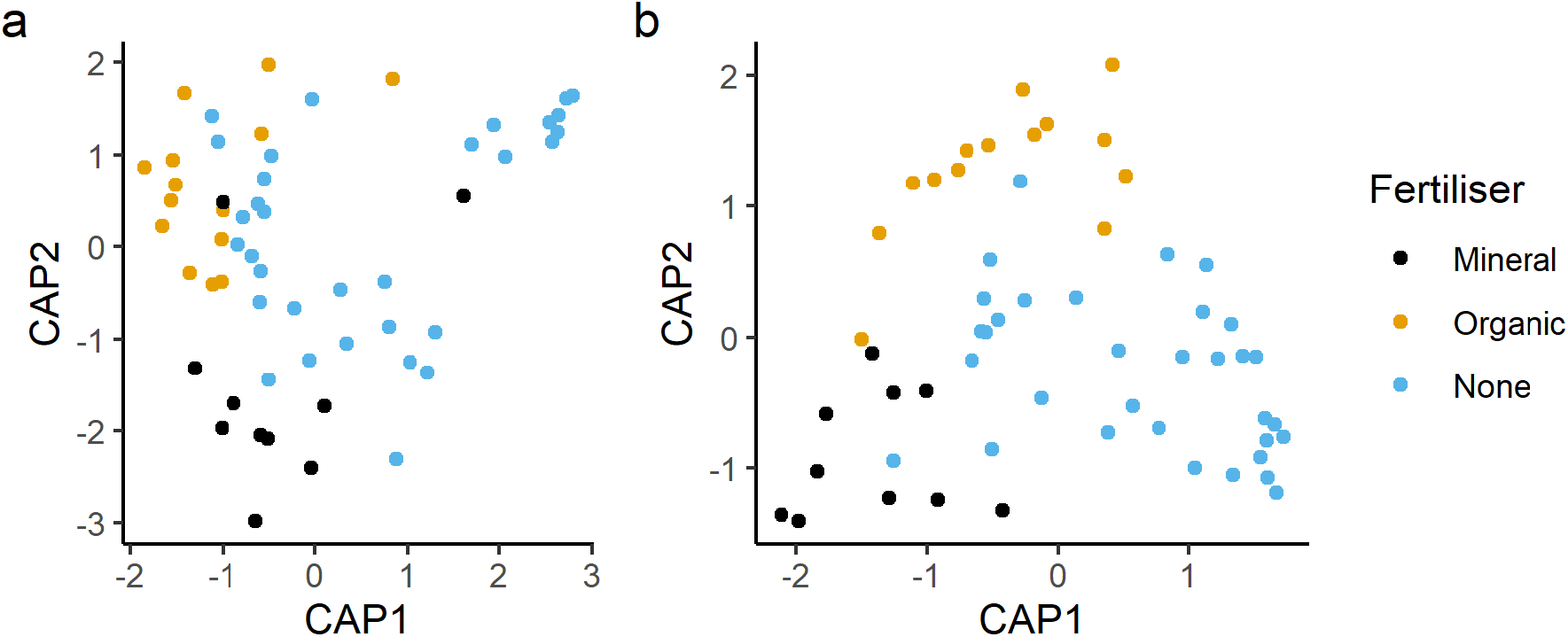
The first two axes of the dbRDA of the bacterial (a) and fungal (b) communities regressed against fertilisation regime. Results were non-significant for bacteria at F_2,46_ = 1.290 and p = 0.102 and significant for fungi at F_2,54_ = 1.238 and p = 0.035.

**Figure 7:**
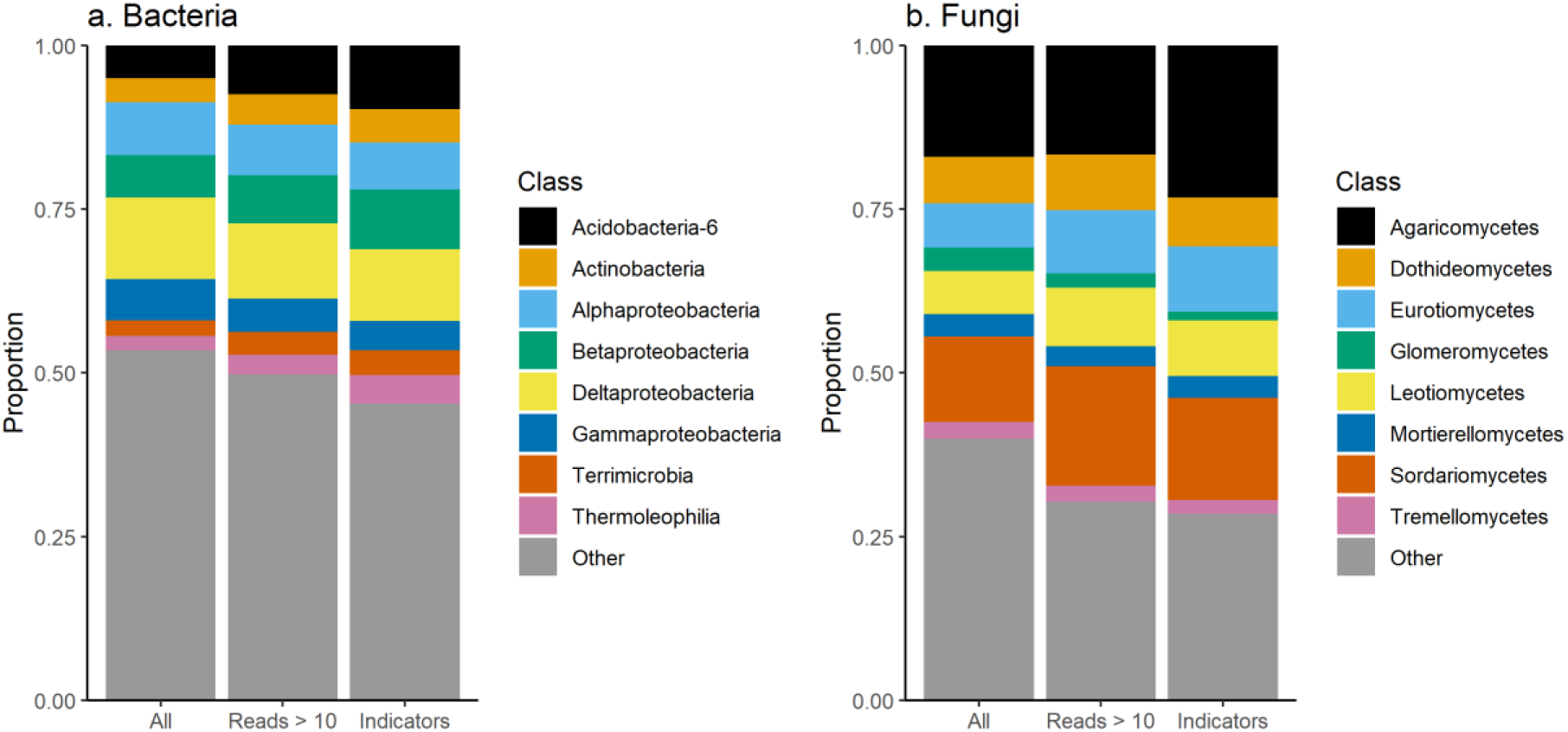
The taxonomic breakdown by class of bacterial (a) and fungal (b) indicators of farm management or soil quality compared to the wider class distribution. The bars represent all bacteria and fungi on the left, followed by all bacteria or fungi with more than 10 reads, and then followed by all bacterial and fungal indicators.

**Table 1:** Bacterial NMDS envfit results, arranged in descending order according to R^2^ value.

| **Variable** | **NMDS1** | **NMDS2** | **R<sup>2</sup>** | **p** |
| --- | --- | --- | --- | --- |
| **pH (CaCl<sub>2</sub>)** | -0.96497 | 0.26238 | 0.8485 | 0.001 |
| **pH (DIW)** | -0.95388 | 0.3002 | 0.8086 | 0.001 |
| **Ellenberg R** | -0.67447 | 0.7383 | 0.4995 | 0.001 |
| **Soil EC** | -0.84651 | 0.53238 | 0.3709 | 0.001 |
| **Total N** | -0.26413 | -0.96449 | 0.3636 | 0.001 |
| **LOI** | -0.19527 | -0.98075 | 0.3533 | 0.001 |
| **Ellenberg F** | 0.2211 | -0.97525 | 0.2756 | 0.002 |
| **Total C** | -0.34766 | -0.93762 | 0.274 | 0.002 |
| **Bulk density** | 0.07565 | 0.99713 | 0.1939 | 0.009 |
| **Stability** | 0.32554 | -0.94553 | 0.1521 | 0.016 |
| **Silt** | 0.06238 | -0.99805 | 0.1518 | 0.019 |
| **Total P** | -0.3447 | -0.93871 | 0.1318 | 0.031 |
| **Plant PCoA axis 3** | -0.62636 | 0.77953 | 0.1241 | 0.033 |
| **Ellenberg N** | -0.72509 | 0.68866 | 0.1143 | 0.055 |
| **Plant PCoA axis 1** | -0.50702 | 0.86193 | 0.111 | 0.055 |
| **Grass cover** | 0.13347 | -0.99105 | 0.107 | 0.067 |
| **Ellenberg L** | -0.21589 | 0.97642 | 0.104 | 0.062 |
| **Clay** | -0.24788 | -0.96879 | 0.0961 | 0.088 |
| **Plant PCoA axis 4** | 0.99688 | 0.07899 | 0.0921 | 0.088 |
| **Weed cover** | -0.21617 | 0.97636 | 0.0805 | 0.105 |
| **Olsen P** | -0.97163 | 0.23653 | 0.0741 | 0.177 |
| **Sand** | -0.02838 | 0.9996 | 0.0699 | 0.172 |
| **Forb richness** | -0.17178 | 0.98514 | 0.0634 | 0.21 |
| **Forb cover** | -0.12679 | 0.99193 | 0.0575 | 0.248 |
| **Lolium perenne cover** | -0.60321 | 0.79758 | 0.0471 | 0.311 |
| **Sheep** | 0.26192 | -0.96509 | 0.0366 | 0.421 |
| **Plant richness** | 0.02657 | 0.99965 | 0.035 | 0.417 |
| **Cattle** | 0.42356 | 0.90587 | 0.0333 | 0.435 |
| **CSM species richness** | 0.73402 | 0.67912 | 0.0331 | 0.452 |
| **Legume richness** | -0.1347 | 0.99089 | 0.0315 | 0.468 |
| **Legume cover** | -0.20849 | 0.97802 | 0.0259 | 0.541 |
| **Plant PCoA axis 2** | -0.23582 | -0.9718 | 0.0158 | 0.668 |
| **Grass richness** | 0.05643 | 0.99841 | 0.0051 | 0.882 |
| **Bare ground cover** | -0.9375 | -0.34798 | 0.0028 | 0.936 |

**Table 2:** Fungal NMDS envfit results, arranged in descending order according to R^2^ value.

| Variable | NMDS1 | NMDS2 | R<sup>2</sup> | p |
| --- | --- | --- | --- | --- |
| pH (CaCl<sub>2</sub>) | -0.97865 | 0.20555 | 0.6514 | 0.001 |
| pH (DIW) | -0.95508 | 0.29635 | 0.6427 | 0.001 |
| Ellenberg R | -0.99466 | -0.10318 | 0.4804 | 0.001 |
| Soil EC | -0.98106 | 0.19373 | 0.315 | 0.001 |
| Ellenberg N | -0.61825 | 0.78598 | 0.3009 | 0.001 |
| Plant PCoA axis 1 | -0.49935 | 0.8664 | 0.2669 | 0.001 |
| Stability | 0.31395 | -0.94944 | 0.2283 | 0.003 |
| Olsen P | -0.48701 | 0.8734 | 0.2055 | 0.008 |
| Legume richness | 0.04607 | -0.99894 | 0.1978 | 0.006 |
| Lolium perenne cover | -0.39181 | 0.92004 | 0.1967 | 0.006 |
| Clay | -0.16572 | -0.98617 | 0.1593 | 0.018 |
| CSM species richness | 0.61437 | -0.78902 | 0.1413 | 0.028 |
| Silt | 0.22583 | -0.97417 | 0.1293 | 0.043 |
| Sand | -0.1286 | 0.9917 | 0.1186 | 0.046 |
| Bare ground cover | -0.08528 | 0.99636 | 0.11 | 0.072 |
| Ellenberg F | 0.50429 | 0.86354 | 0.1023 | 0.065 |
| Ellenberg L | -0.419 | -0.90799 | 0.0825 | 0.126 |
| Plant PCoA axis 4 | 0.99975 | 0.02238 | 0.0754 | 0.15 |
| Grass richness | 0.2109 | -0.97751 | 0.0654 | 0.199 |
| Total P | -0.99999 | 0.00523 | 0.0632 | 0.21 |
| Weed cover | -0.3488 | 0.9372 | 0.0625 | 0.206 |
| Plant richness | 0.35284 | -0.93568 | 0.0612 | 0.225 |
| Total N | -0.8692 | -0.49447 | 0.0591 | 0.237 |
| Total C | -0.99944 | -0.03346 | 0.0551 | 0.273 |
| Plant PCoA axis 3 | -0.99738 | 0.07234 | 0.0502 | 0.275 |
| Forb cover | -0.13627 | -0.99067 | 0.0487 | 0.288 |
| Forb richness | 0.0322 | -0.99948 | 0.039 | 0.392 |
| Plant PCoA axis 2 | -0.44484 | -0.89561 | 0.0356 | 0.448 |
| Sheep | 0.85609 | -0.51682 | 0.031 | 0.476 |
| Bulk density | -0.06753 | 0.99772 | 0.0255 | 0.531 |
| Cattle | 0.34364 | -0.9391 | 0.0238 | 0.529 |
| LOI | -0.82729 | -0.56177 | 0.0219 | 0.588 |
| Legume cover | -0.24543 | -0.96941 | 0.0137 | 0.71 |
| Grass cover | 0.814 | 0.58087 | 0.011 | 0.742 |

